# A single injection of anti-PD-L1 blocking antibody induces a transient reduction in tau pathology in P301S (PS19) mouse model of tauopathy

**DOI:** 10.1101/2025.09.28.679024

**Authors:** Alexander Kertser, Kuti Baruch, Michal Schwartz

## Abstract

Blocking the inhibitory PD-1/PD-L1 immune checkpoint pathway has been shown to arrest cognitive decline and reduce multiple aspects of brain pathology in various mouse models of amyloidosis and tauopathy. In this study, we evaluated whether anti-PD-L1 treatment would be effective in a tauopathy model characterized by rapidly progressing pathology. Using the P301S (PS19) transgenic mouse model, we found that a single injection of anti-PD-L1 antibody significantly attenuated cognitive deficits, with effects detectable 1-month post-treatment, aligned with the kinetics reported in other models. This cognitive benefit was observed irrespective of microglial TREM2 signalling. The effect on brain pathology was transient; phosphorylated tau in the brain, and total tau in the cerebrospinal fluid were significantly reduced 14 days post-treatment but not at 28 days. These findings indicate that PD-L1 blockade is effective across multiple disease models, while emphasizing the importance of carefully monitoring the timing of effect assessments, treatment outcomes, and dosing frequency, especially in models with accelerated disease progression.

## Introduction

Tauopathies comprise a heterogeneous group of neurodegenerative disorders, characterized by accumulation of abnormally modified tau proteins that lead to synaptic dysfunction, local inflammation, progressive neuronal loss, and cognitive and/or motor decline. Among preclinical mouse models, the PS19 transgenic line (human P301S mutation 1N4R tau under the prion promoter) is widely used, because it develops robust cortical and hippocampal tau pathology, pronounced neuronal loss, and exhibits an aggressive clinical course with early mortality (Yoshiyama et al., 2007; Zhong et al., 2024). This rapid trajectory provides a stringent setting to test disease-modifying interventions other than those that are directed to the tau pathology and poses the question of whether therapies that show benefit in slower experimental models can overcome the rapid disease kinetics of the PS19 model. This challenge is underscored by the fact that, while the accumulation of misfolded proteins in the central nervous system (CNS) is the hallmark of tauopathy mouse models, clinical trials targeting these proteins for removal with anti-tau antibodies have so far demonstrated limited therapeutic impact (Imbimbo et al., 2023). Such limited efficacy likely reflects the multifactorial nature of CNS pathologies, in which protein aggregation coexists with other contributing mechanisms - most notably, chronic neuroinflammation (Leng & Edison, 2021; Wong-Guerra et al., 2023) .

Given this, attention has increasingly shifted towards the peripheral immune system, which is now recognized as a key factor in CNS protection and repair (Castellani et al., 2023; Schwartz & Colaiuta, 2024; Kim & Kipnis, 2025). In mouse models of Alzheimer’s disease (AD) and tauopathy, inducing a peripheral immune response in a well-controlled manner was shown to modulate blood-cerebrospinal fluid (CSF) barrier function to facilitate homing of specialized immune cells to the CNS, and attenuate disease progression (Baruch et al., 2015; Rosenzweig et al., 2019). Conversely, peripheral immune suppression or deficiency has been identified as a detrimental factor that further compromises brain support in aging and neurodegenerative conditions (Schwartz & Kipnis, 2011; Marsh et al., 2016). Consequently, therapeutic strategies aimed at restoring the crosstalk between the peripheral immune system and the brain may offer an indirect yet potent avenue to mitigate CNS disease progression, even if these approaches do not prevent its onset.

Targeting inhibitory immune checkpoint pathways, such as the programmed cell death-1 (PD-1)/PD-L1 (Baruch et al., 2016; Rosenzweig et al., 2019; Ben-Yehuda et al., 2021; Xing et al., 2021; Zou et al., 2021; Liu et al., 2021; Dvir-Szternfeld et al., 2022) or TIM-3 (Kimura et al., 2025), has emerged as a new immune-mediated therapeutic strategy in the context of neurodegenerative diseases. In animal models of amyloidosis, and in the DM-hTau (K257T/P301S) mouse model of tauopathy, it was demonstrated that a single injection targeting either the PD-1 receptor or its ligand, PD-L1, is sufficient to evoke an immune response that culminates in beneficial functional effects (Baruch et al., 2016; Rosenzweig et al., 2019).

Here, we assessed whether a single injection of anti-PD-L1 blocking antibody, administered in a dosing previously shown to be effective in slower-progressing disease models of AD and tauopathy, could be effective in PS19 mice. Furthermore, we also tested whether the PD-L1-induced benefits are dependent on TREM2 function, given that rare TREM2 variants confer strong genetic risk for AD and have been previously linked to impaired microglial responses in neurodegeneration (Leyns et al., 2017). We found that in PS19 mice a single injection of anti-PD-L1 yielded a beneficial effect on cognitive performance 28 days post-injection, consistent with the observed effect characterized in previously tested models. However, the impact on disease pathology was only transient, preceding the observed cognitive improvement. These findings suggest that the rate of disease progression, at least in experimental models, should be taken into consideration when testing immunotherapy.

## Materials and Methods

### Animals

B6;C3-Tg(Prnp-MAPT*P301S)PS19Vle/J mice (PS19; Jackson Laboratory, stock no. 002726) were bred and maintained at the Weizmann Institute of Science (WIS) animal facilities. PS19 mice were cross-bred with Trem2 knockout (Trem2^−/−^; TREM2 KO) mice, originally obtained from the laboratory of Marco Colonna (Washington University). In one of the studies, PS19 mice were transferred as adults to the animal facilities of Science in Action Ltd., Israel, where they received treatment and were sacrificed for tissue collection. Genotyping was performed by PCR analysis of tail DNA. Wild-type controls in each experiment were non-transgenic littermates from the relevant mouse colonies. Both male and female mice were used in the studies and, when possible, were equally distributed between experimental groups. All animal experiments were approved by the Institutional Animal Care and Use Committee (IACUC) of the Weizmann Institute of Science.

### Antibodies injections

Two anti–PD-L1 antibodies were used in this study. The first was a surrogate of IBC-Ab002, consisting of a human anti-mouse PD-L1 antibody on an IgG1 backbone engineered with Fc modifications to abrogate effector function and enhance clearance from the circulation. The second was a rat anti-mouse PD-L1 monoclonal antibody (clone 10F.9G2, IgG2b isotype; Bio X Cell). Antibodies were diluted in sterile phosphate-buffered saline (PBS) and administered intraperitoneally (i.p.) as a single bolus injection (200 µL; 1.5 mg per mouse). Matching isotype control antibodies (rat IgG2b, clone LTF-2, Bio X Cell, or human IgG1 anti-B12) were used in parallel.

### Flow cytometry

Fresh whole blood was processed using ammonium–chloride–potassium (ACK) lysing buffer to remove erythrocytes. Remaining cells were washed, passed through a 70-µm nylon mesh, and stained with fluorochrome-conjugated antibodies against surface markers: FITC anti-CD62L (clone MEL-14; BioLegend, cat. #104406), PE anti-CD4 (clone GK1.5; BioLegend, cat. #100408), APC anti-CD44 (clone IM7; BioLegend, cat. #103012), APC/Cy7 anti-PD-1 (clone 29F.1A12; BioLegend, cat. #135224), and FITC anti-CD3 (clone 145-2C11; BioLegend, cat. #100306). Data were acquired on a CytoFLEX flow cytometer (Beckman Coulter) and analyzed using FlowJo software (BD Biosciences).

### Cognitive assessments

Mice underwent daily 3-min handling sessions for five consecutive days prior to behavioural testing. All behavioural experiments were performed by investigators blinded to the treatment allocation.

### T-maze cognitive task

Spatial short-term memory was assessed using the T-maze, which exploits the natural tendency of mice to explore novel environments. The maze consisted of a 57-cm central alley and two 45-cm arms extending at right angles, each 10 cm wide with 10-cm high walls. The task consisted of two trials separated by a 5-min inter-trial interval, during which mice were placed in individual holding cages. In the acquisition trial (8 min), each mouse was introduced at the center of the maze and allowed to explore two accessible arms, while the third arm was blocked. In the retention trial (3 min), all three arms were opened, and the mouse was allowed to freely explore. Exploration time was calculated as the total time spent in the novel arm, which reflects the animal’s ability to recognize and preferentially explore an unfamiliar environment.

### Novel Object Recognition cognitive task

Recognition memory was assessed using the novel object recognition (NOR) test. The apparatus consisted of a gray plastic arena measuring 45 × 45 × 50 cm, with visual cues on the walls. The task spanned 2 days. On the first day, mice were habituated to the empty arena for 20 minutes. On the second day, mice underwent a 10-minute familiarization trial with two identical objects. Following a one-hour interval, a 6-minute test trial was conducted in which one of the familiar objects was replaced with a novel object. A percentage of exploration time was calculated as the ratio of time spent interacting with the novel object to the total time spent interacting with both objects, providing an index of recognition memory.

### Terminal tissue collection

Cerebrospinal fluid (CSF) was collected by cisterna magna puncture. Mice were then transcardially perfused, and brain tissues were extracted and stored at -80 °C until tissue analysis.

### Biochemical assays for tau species quantification

Cortices and hippocampi were homogenized on ice using a Dounce homogenizer in lysis buffer (5 µL per 1 mg tissue; 1% Triton X-100, Halt™ protease inhibitor cocktail [1:100; Thermo Scientific] in phosphate-buffered saline). Protein concentrations were determined using the BCA protein assay kit (Pierce) according to the manufacturer’s instructions, and samples were analyzed at three dilutions ranging from 1 to 0.11 mg/mL protein. CSF samples were diluted 1:8 in assay buffer. Homogeneous time-resolved fluorescence (HTRF) kits for MAPT, phospho-S404, and phospho-S202/T205 detection (Revvity, formerly Cisbio) were used according to the manufacturer’s instructions. Readings were obtained using a Spark multimode plate reader (Tecan).

### Neurofilament light chain (NfL) quantification

NfL levels were measured in cerebrospinal fluid (CSF) using the NfL ELISA kit (Uman Diagnostics) according to the manufacturer’s instructions, with minor modifications. CSF samples were diluted 1:50, incubated overnight with agitation (rather than for 1 hour), and the standard curve was extended at the lower range by two additional serial dilutions. Serum NfL was quantified using the human NfL kit on the SIMOA platform (Quanterix) according to the manufacturer’s instructions.

### Statistics

Statistical analyses were performed using GraphPad Prism software (version 10; GraphPad Software, San Diego, CA). Data are presented as mean ± standard error of the mean (SEM). Comparisons between two groups were performed using unpaired two-tailed Student’s t-test. For comparisons involving more than two groups, one-way or two-way analysis of variance (ANOVA) with appropriate post hoc multiple comparison tests was applied. In all analyses, a p-value < 0.05 was considered statistically significant. The number of biological replicates (n) and the statistical test used are indicated in the figure legends.

## Results

### Effect of PD-L1 blockade on cognitive performance in PS19 mice

We first assessed the peripheral immune response triggered by administration of an antibody directed against PD-L1 (anti-PD-L1) to PS19 transgenic mice. Male and female PS19 transgenic mice received intraperitoneal injections of either a sub-therapeutic dose (0.1 mg/mouse) or an effective dose (1.5 mg/mouse) of PD-L1 blocking antibody engineered to exhibit rapid clearance from the circulation (T1/2 in mice = 14.5h) (Baruch et al., 2020). A single injection of the effective dose (1.5 mg/mouse) of anti-PD-L1 led to an increase in frequencies of circulating memory (CD44^+^) CD4^+^ T cells expressing PD-1 (**Figure 1a**), a cell surface marker of immune activation, indicating that the immune system of PS19 mice was responsive to treatment. These results also confirmed that the effective dose of anti-PD-L1 needed for triggering a peripheral immune response in PS19 mice is similar to the dose previously identified in other mouse models of AD and tauopathies (Rosenzweig et al., 2019).

**Figure 1.**
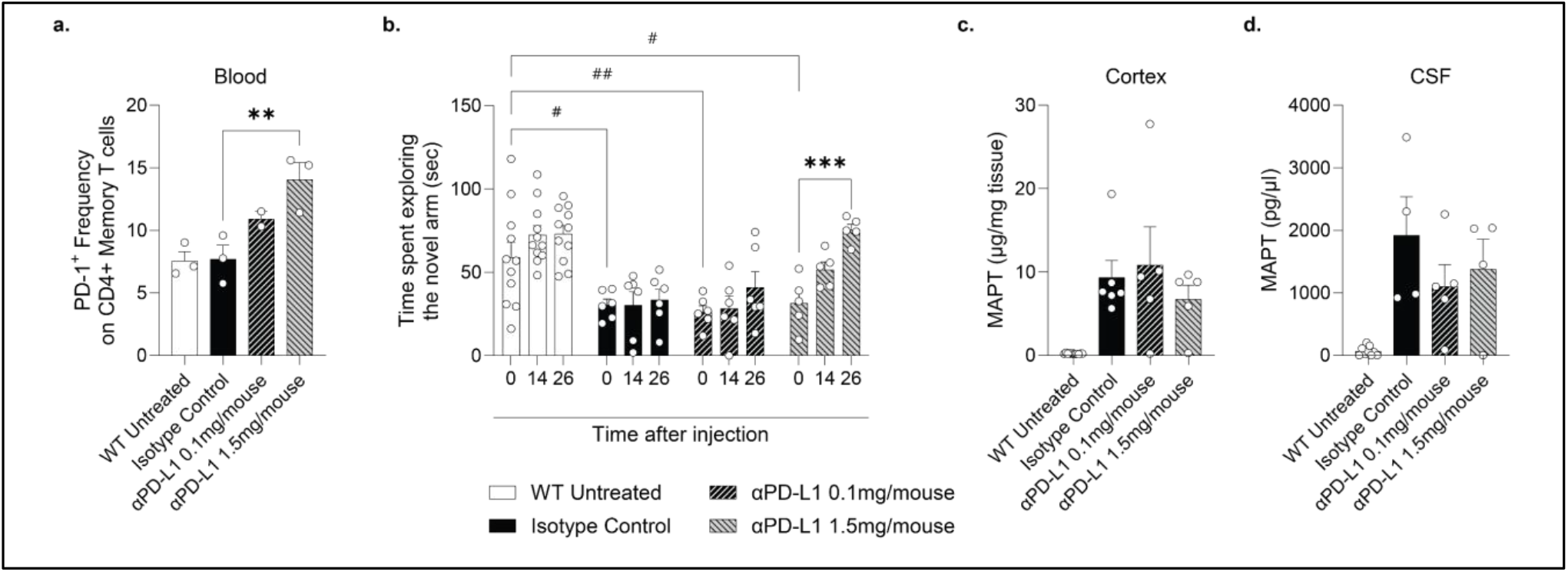
A single injection of anti-PD-L1 to PS19 mice leads to arrest of cognitive loss at 4 weeks post-treatment without direct effect on the level of Tau protein. 7–8-month-old PS19 mice injected with either 0.1 or 1.5mg/mouse anti-PD-L1 antibody (short half-life variant) were tested for (a) PD-1 expression on blood-borne memory (CD44^+^) CD4 T cells on day 3 after injection. Isotype control injected mice and non-transgenic (WT) untreated mice served as controls. n=2-3 per group; one-way ANOVA followed by Fischer post-hoc test; error bars represent mean ± SEM; **, p< 0.01 between indicated groups. In a separate group of mice, (b) time spent in the novel arm of a T-maze was determined at baseline (8 months old; before drug administration) and on days 14 and 26 following injection. Total Tau (MAPT) levels were calculated using HTRF in (c) cortex and (d) CSF on day 28 after treatment. n=5-11 per group; two-way ANOVA followed by Dunnett’s multiple comparisons test; error bars represent mean ± SEM; **, p< 0.01 between indicated groups.

PS19 mice display deficits in cortex-dependent working memory and exploration behaviour (Ahmad et al., 2021). We therefore assessed cognitive performance using the T-maze task, in which mice are exposed to a novel environment in a T-shaped two arm maze, and scored according to the time spent exploring its arms. At 8 months of age, PS19 mice exhibited a pronounced reduction in exploration behaviour relative to their non-transgenic wild-type littermates, across all timepoints tested (**Figure 1b**). Following anti-PD-L1 treatment (1.5mg/mouse), the mice showed better exploration behaviour than mice treated with an isotype control antibody at day 14, with an effect becoming significant by day 26 following the treatment. Mice that received a sub-therapeutic antibody dose (0.1 mg/mouse), showed no difference in cognitive performance relative to control-treated mice (**Figure 1b**).

Mice were euthanized 28 days after treatment for terminal tissue collection. Cortical brain tissue and CSF samples were collected and analysed for levels of microtubule-associated protein tau (MAPT) using a Homogeneous Time-Resolved Fluorescence (HTRF) assay. As expected, PS19 mice exhibited high expression of human MAPT in both cortex tissues and CSF (**Figure 1c,d**). We found that at this timepoint, in which we observed functional effect of anti-PD-L1 treatment on behaviour, no effect on MAPT brain pathology was observed in the anti-PD-L1-treated group of mice compared with the isotype control treated group (**Figure 1c,d**).

### PD-L1 blockade in the PS19 mouse model leads to a detectable yet temporary reduction in brain Tau S404 phosphorylation

Intrigued by the improvement in cognitive performance in the absence of detectable changes in total MAPT at the same timepoint after treatment, we hypothesized that effects on tau pathology might occur earlier and precede the functional behavioural response. To test this, 6-month-old PS19 mice were treated with the effective dose of the anti-mPD-L1, and brain tissues were collected for terminal analysis on either day 14 or 28 post-treatment. Tau hyperphosphorylation at residue S404, a site linked to aggregated, insoluble Tau, was measured by HTRF in the cortex and hippocampus brain tissues. The mice treated with anti-PD-L1–exhibited a significant reduction in cortical S404 phosphorylation only at day 14, with no effect on day 28 following a single injection relative to the levels in the isotype control treatment group (**Figure 2a**). Notably, unlike the cortex tissue, no effects were observed in the hippocampus at any time points tested (**Figure 2b**). We also examined neurofilament light chain (NfL) levels in CSF and serum, as a biomarker of neurodegeneration and disease progression (Bacioglu et al., 2016). Despite being age-matched at the time of the treatment, a wide variation in NfL levels was observed, spanning several orders of magnitude within the same group in both CSF and serum. Therefore, although anti-PD-L1–treated mice exhibited a trend toward lower NfL levels compared to isotype controls, the differences did not reach statistical significance at either day 14 or day 28 following a single injection (**Figure 2c,d**).

**Figure 2.**
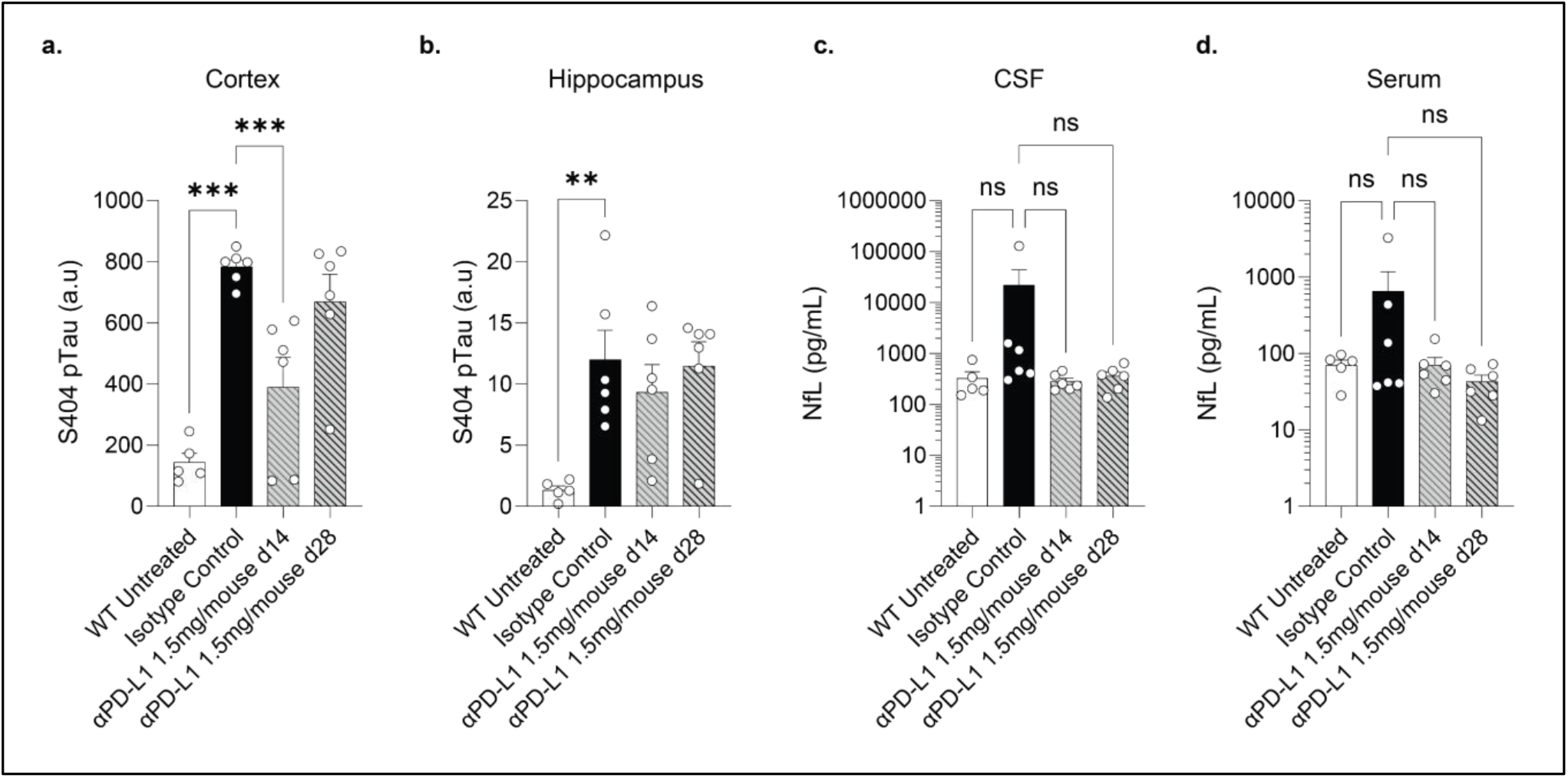
PD-L1 blockade in PS19 mouse model leads to a detectable yet temporary reduction in brain Tau S404 phosphorylation. PS19 mice received either 1.5mg/mouse anti-PD-L1 blocking antibody or isotype control at 5-6 months of age. Non-transgenic (WT) littermates were used as a negative control. (a) Cortical and (b) hippocampal levels of S404 hyperphosphorylated Tau protein were measured using HTRF on days 14 and 28 following injection. (c) CSF and (d) serum levels of NfL were measured using ELISA or SIMOA, respectively. n=5-6 per group; one-way ANOVA followed by Fischer post-hoc test; error bars represent mean ± SEM; **, p< 0.01; ***, p< 0.001; ns = non-significant, between indicated groups.

### Effect of PD-L1 blockade on different tau species

Given the transient effects observed on brain pathology in 6-month-old mice following anti-PD-L1 treatment, we decided to repeat this experiment in a larger cohort of older (9-month-old) PS19 mice, this time using a different anti-PD-L1 blocking antibody (Rat IgG2b anti-mouse-PD-L1; clone 10F.9G2). The treatment effect on cortical tau pathology was assessed at two sites of tau phosphorylation: S404, which we previously measured, and S202/T205, which is associated with early pre-tangle pathology. At day 14 following treatment with the effective antibody dose (1.5 mg/mouse), we observed reduced levels of S404 phosphorylation, with no effect on the levels of S202/T205 relative to isotype-treated controls (**Figure 3a**). In the CSF compartment, anti-PD-L1 treatment led to a reduction in total MAPT levels at day 14 (**Figure 3b**). NfL levels showed wide variability across mice and did not show statistical significance (**Figure 3c**). These data highlight that a single injection of anti-PD-L1 antibody transiently modified certain pathological readouts in the brain of PS19 mice.

**Figure 3.**
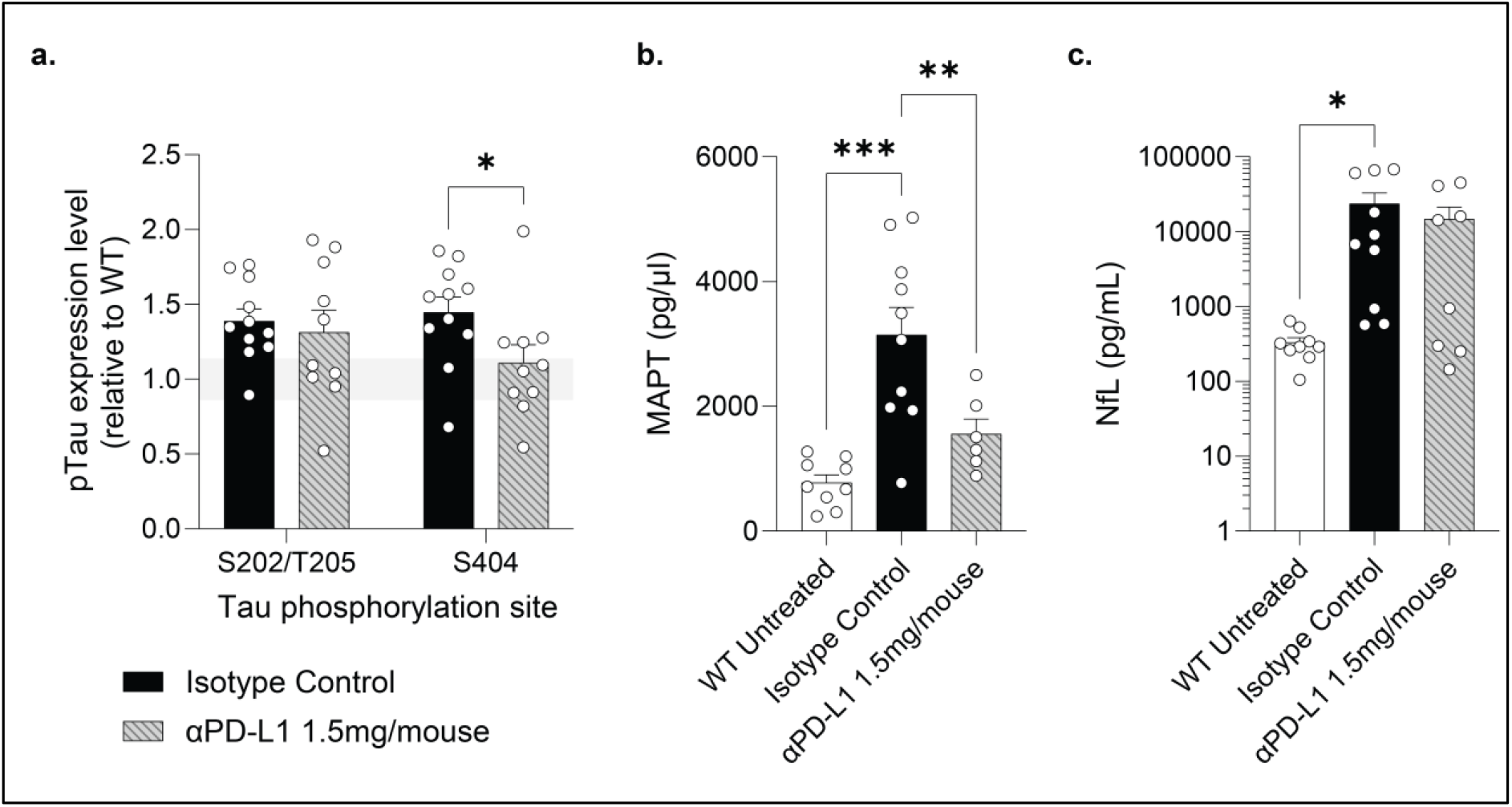
Effect of PD-L1 blockade on different tau species. Male and female 9-month-old PS19 mice were treated with either 1.5mg/mouse anti-PD-L1 blocking antibody or isotype control. Non-transgenic (WT) littermates were used as a negative control. (a) Levels of Tau protein phosphorylation on S202/T205 or S404 measured in the brain cortices of male and female PS19 (Tg) mice 14 days after treatment. Grey area shows standard error margins of levels measured in WT untreated mice. CSF levels of (b) total Tau (MAPT) and (c) NfL were measured on day 14 following injection in the same experiment. n=8-11 per group; one-way ANOVA followed by Fischer post-hoc test; error bars represent mean ± SEM; *, p< 0.05; **, p< 0.01; ***, p< 0.001 between indicated groups.

### PD-L1 blockade reduces cognitive deficit in PS19 mice independently of TREM2

TREM2 is functionally linked with microglial activation (Keren-Shaul et al., 2017) and senescent microglia (Rachmian et al., 2024). Moreover, complete deficiency has been reported to attenuate neurodegeneration in PS19 mice by damping microglial activation (Leyns et al., 2017), whereas partial loss can exacerbate tau pathology and inflammation (Sayed et al., 2018). Importantly, checkpoint blockade has shown efficacy in an amyloid model even in the absence of TREM2 (Dvir-Szternfeld et al., 2022). To test whether the observed beneficial effect of PD-L1 blockade in PS19 is dependent on TREM2 signalling, we generated a line of PS19 mice lacking TREM2, and evaluated the response to treatment.

PS19/Trem2^−/−^ mice treated with anti-PD-L1 (1.5 mg/mouse) at 8 months of age were tested 28 days following antibody injection. Treated mice spent more time exploring the novel arm of the T-maze (**Figure 4a**) and showed increased interaction with a novel object in the recognition task (**Figure 4b**), indicating improved cognitive performance. As in TREM2^+/+^ mice, no effects on S404 tau phosphorylation were detected at this late time point (**Figure 4c**). To exclude the possibility that these findings reflected slower disease progression in TREM2-deficient PS19 mice, we tested an additional cohort of mice injected at a slightly older age (9 months of age), when cognitive deficits are more pronounced. At this disease stage, as well, PD-L1 blockade led to an improvement of exploration behaviour, which was comparable to that of their non-transgenic littermates (**Figure 4d**). These results suggest that the cognitive benefits of PD-L1 blockade are preserved in the absence of TREM2, supporting a mechanism of action independent of TREM2 signalling.

**Figure 4.**
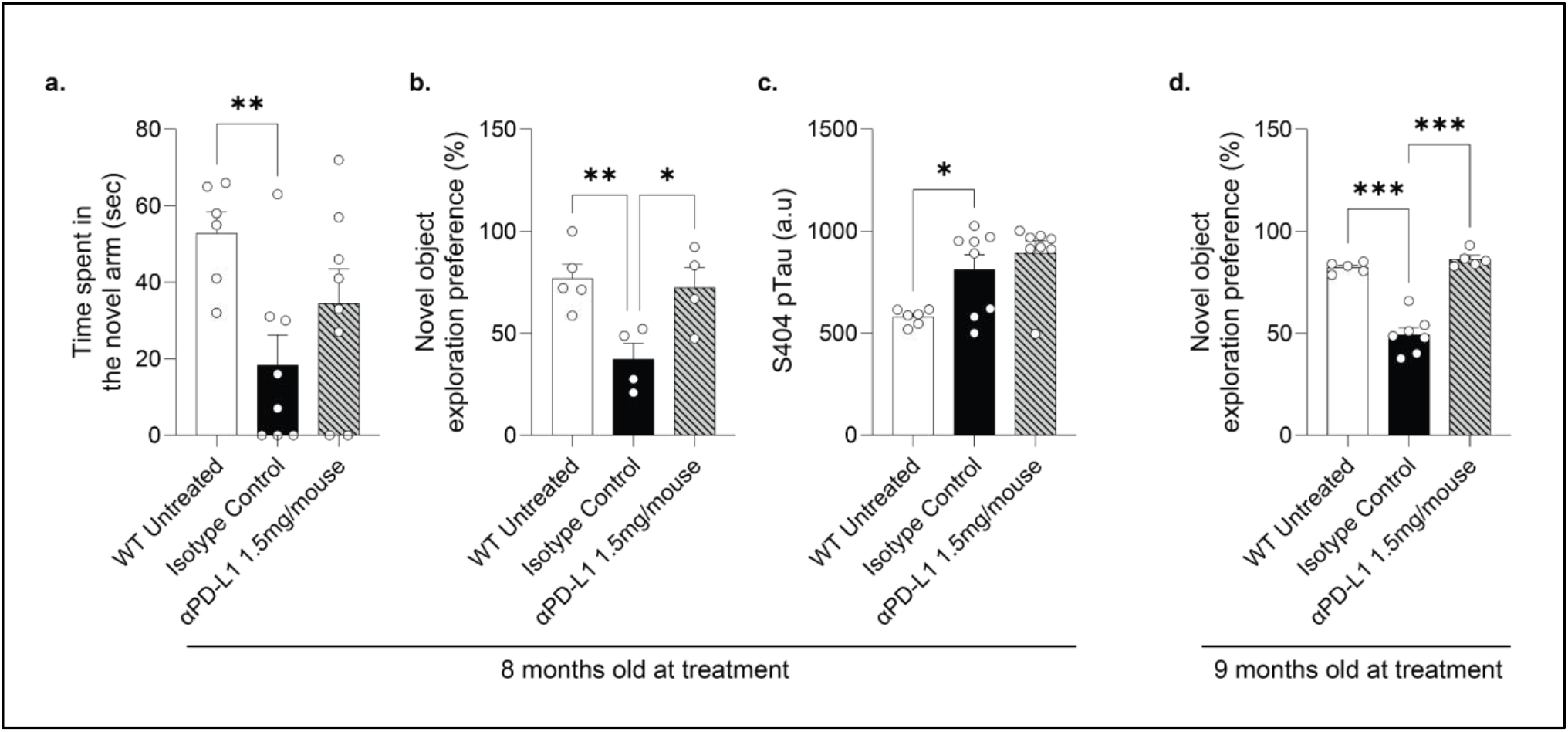
PD-L1 blockade improves cognitive performance in PS19 mice independently of TREM2. PS19 mice were cross-bred with Trem2^-/-^ (knock-out (KO)) mice. Isotype control or anti-PD-L1 (1.5mg/mouse) antibody were administered to KO mice at the age of 7 months, and cognitive performance was tested 28 days later in (a) T-maze exploration, and (b) novel object recognition tasks. (c) Cortical levels of S404 Tau phosphorylation were measured using HTRF assay. n=6-8 per group; one-way ANOVA followed by Fischer post-hoc test; error bars represent mean ± SEM; *, p< 0.05; **, p< 0.01; between indicated groups. (d) A separate group of PS19 mice on a Trem2^-/-^ background were treated with anti-PD-L1 at the age of 8-9 months and tested for changes in cognitive performance using the novel object recognition task. n=5-6 per group; one-way ANOVA followed by Fischer post-hoc test; error bars represent mean ± SEM; ***, p< 0.001; between indicated groups.

## Discussion

This study aimed to test whether a single injection of PD-L1 blocking antibody, a therapeutic approach demonstrated in several mouse models of AD and tauopathies, can affect behaviour and brain pathology readouts in the aggressive PS19 (P301S) tauopathy model. First, we found that a single effective dose of anti-PD-L1 in PS19 mice rapidly engages the peripheral immune system (increased PD-1 on circulating memory CD4^+^ T cells), which was accompanied by significant cognitive benefits at the timepoint of ∼4 weeks after treatment. Second, the impact on brain pathology was found to be time- and tau species-specific. We detected a transient reduction in cortical tau S404 phosphorylation alongside a decrease in CSF total tau at 14 days post single-injection, but not at 28 days after treatment. Lastly, we found that the beneficial effect on behaviour was also observed in the absence of TREM2, indicating that PD-1/PD-L1 pathway blockade did not depend on TREM2 signalling.

Our results show that transient beneficial effect in reducing brain pathology can be elicited by a single administration of antibody blocking the PD-1/PD-L1 pathway in PS19 mice. Yet, in contrast to other tested mouse models such as 5xFAD and DM-hTau (Rosenzweig et al., 2019), in the PS19 line this effect was detected on day 14, but not on day 28 after a single injection. It is possible that the beneficial effects on pathology are outpaced by the intrinsic aggressiveness of the model, explaining why effects on proteinopathy were found to be transient. Alternatively, it is possible that the treatment’s benefit across all models is primarily driven by the dampening of brain inflammation, with all other positive outcomes downstream of that effect. Accordingly, sustained therapeutic effect over time may reflect enhanced neuronal survival rather than a reduction in proteinopathy. Aligned with these findings, Chen et al. tested chronic treatment with anti-PD-1 antibody in TE4 (P301S/E4) mouse model of tauopathy, beginning at 8 months of age, and demonstrated a significant beneficial effect in reducing tau-mediated neurodegeneration and brain atrophy (Chen et al., 2023).

Harnessing the peripheral immune system for brain support is fundamentally different approach from any therapies that directly target brain proteinopathy. In mouse models of tauopathies, several anti-tau therapies (passive immunization with antibodies against phospho-tau epitopes, or vectorized anti-tau fragments) have shown direct and sometimes robust effects on insoluble tau burden, neurodegeneration, and behaviour (Ising et al., 2017; Sankaranarayanan et al., 2015; Yanamandra et al., 2015). In contrast, when a therapy acts indirectly on proteinopathy, as in the case of anti-PD-L1 administration, its impact on disease progression is mediated through immune-cell recruitment to the CNS, affecting primarily the neuroinflammatory miliue. Indeed, in the relatively slowly-progressing tauopathy model of DM-hTau (K257T/P301S under the native MAPT promoter) mice, the treatment effect of anti-PD-L1 antibody on neuroinflammation was followed by an effect on tau pathology, observed 1 month after a single injection (Rosenzweig et al., 2019; Ben-Yehuda et al., 2021). Mechanistically, the observed effect in DM-hTau mice was found to be associated with recruitment of both monocyte-derived macrophages and Foxp3^+^ regulatory T cells to the brain (Ben-Yehuda et al., 2021). Similarly, increased Foxp3^+^ regulatory T cells in the brain were associated with a beneficial effect on brain atrophy in the TE4 tauopathy mouse model (Chen et al., 2023).

The behavioural effects in PS19 mice described here are consistent with prior reports demonstrating that a single injection of antibody blocking the PD-1/PD-L1 checkpoint pathway, induced effects emerging well after the initial immune cell activation observed in the periphery (Baruch et al., 2016; Rosenzweig et al., 2019; Ben-Yehuda et al., 2021). Dose dependency in the current study (demonstrating benefit at 1.5 mg/mouse, with no effect with 0.1 mg/mouse or isotype control antibody) also echoes previous published dose-response studies that were demonstrated in the 5xFAD mouse model of amyloidosis and DM-hTau mouse model of tauopathy (Rosenzweig et al., 2019). Together, these data support a threshold requirement for checkpoint engagement to trigger the desired immune response that ultimately leads to brain functional benefit.

Investigating brain pathology, we found a time-limited (at day 14) reduction in cortical tau S404-but not S202/T205-phosphorylation; no effects were observed in the hippocampus at this time point. This pattern fits a well-described epitope hierarchy in which AT8 (pS202/pT205) marks earlier phosphorylation events, while PHF-1 (pS396/pS404) tracks later, aggregation-prone conformations enriched in insoluble fractions; thus, a transient decrease in PHF-1 with stable AT8 is compatible with a transient shift in the aggregated pool, rather than broad reduced phosphorylation across sites (Neddens et al., 2018; Wegmann et al., 2021; Cantrelle et al., 2021). The accompanying transient reduction in total CSF tau strengthens this interpretation, as CSF t-tau is considered as a sensitive reporter of tau production/turnover and is modifiable by therapies that lower tau load (e.g., MAPT-targeting ASO BIIB080), even when gross histopathology has not yet diverged (Edwards et al., 2023). Brain-region specificity is also consistent with prior studies of PS19, showing brain-region differences in synaptic remodelling and tau compartmentalization, in which hippocampus and cortical regions can diverge in synaptic tau dynamics and structural plasticity in cohorts of similar age (Walker et al., 2021).

The behavioural benefit following PD-L1 blockade was observed also in PS19/Trem2^−/−^ mice, indicating that it does not require TREM2 signalling. This is consistent with evidence from the 5xFAD mouse model of amyloidosis, showing that bone-marrow–derived macrophages can mediate disease modification via TREM2-independent pathways (Dvir-Szternfeld et al., 2022).

We note that not all reported studies, using models of AD and tauopathy, showed a cognitive benefit following PD-1/PD-L1 blockade; for example, chronic weekly anti-PD-1 dosing for 12 weeks in JNPL3 mice did not alter cognitive or tau endpoints (Lin et al., 2019). Such discrepancies likely reflect differences in model background, disease stage at intervention, and, critically – the regimen (prolonged weekly dosing vs. intermittent “pulses” with delayed readouts), emphasizing the importance of schedule and timing for periphery-to-brain mechanisms (Baruch & Yoles, 2020).

Several limitations of this study should be acknowledged. Our biochemical analysis of brain tissue focused on pS404 and pS202/T205 at limited timepoints post-treatment; additional temporal profiling would better define onset and durability of treatment effect. In addition, our behavioural assays focused on exploration and recognition tasks; inclusion of additional cognitive and motor paradigms might capture domain-specific sensitivity. Further studies are required to determine whether the cognitive effects in PS19 mice are primarily driven by modulation of neuroinflammation, with other downstream effects being transient, and require repeated dosing for sustained effect. Notably, we tested a single pulse regimen; exploring repeated dosing in PS19 could extend benefit, yet long term effect may be limited by the aggressiveness of this model.

